# Social Novelty Recruits a Dysfunctional Nucleus Accumbens Ensemble That Drives Social Avoidance in a *Shank3^−/−^* Autism Model

**DOI:** 10.64898/2026.06.04.730003

**Authors:** Oakleigh M. Folkes, Meaghan Donahue, Rafael E. Perez, Angélica Minier-Toribio, Paola N. Negrón-Moreno, Marina R. Picciotto, Yong-Hui Jiang

**Author notes:** **Correspondence:** Oakleigh Folkes, Ph.D. Yong-hui Jiang, MD, PhD.

## Abstract

Social behavior deficits are a common symptom of neuropsychiatric disorders, including autism spectrum disorder (ASD), but there are limited pharmacological treatments for these symptoms. Understanding how neurons encode social information will give insight into identifying novel pharmacological targets to address this unmet need.

*SHANK3* encodes a postsynaptic scaffold protein and is a common risk gene for several neuropsychiatric disorders characterized by social deficits, including ASD. Our lab previously developed the *Shank3*^Δ*e4–22*^ mouse model, which shows a loss of social preference and altered connectivity in the nucleus accumbens (NAc), a critical region for social behaviors. However, it remains unknown how *Shank3* deletion alters the encoding of social cues in the NAc.

To address this gap in knowledge, we characterized the function of neurons activated by social interaction, or social ensembles, in WT and *Shank3*^Δ*e4–22*^ mice using a combination of genetic capture techniques, chemogenetics, optogenetics, and one-photon calcium imaging. We show that NAc social ensembles of WTs and *Shank3*^Δ*e4–22*^ mice drive opposing social behaviors: while NAc social ensembles encode appetitive social cues in WT mice, they encode avoidance in *Shank3*^Δ*e4–22*^ mice. We further find that, in *Shank3*^Δ*e4–22*^ mice, NAc neurons are hyperactive and have hypermodulatory responses to social novelty. Suppressing the activity of social ensembles during social novelty prevents future social avoidance and restores social investigation in *Shank3*^Δ*e4–22*^ mice. Taken together, our data show that *Shank3*^Δ*e4–22*^ mice have an enhanced NAc response to social novelty that actively drives social aversion, rather than a loss of social motivation.

## Introduction

Social behavior encompasses a broad range of domains, including appetitive social approach and aversive social avoidance.^1^ Positive social interactions and social support are well-established protective factors against the worsening and development of many disorders, whereas social withdrawal and isolation can induce chronic health conditions like depression and anxiety.^2–7^ In neuropsychiatric disorders, including Autism Spectrum Disorder (ASD), impairments in social communication and interaction substantially limit positive social behavior and increase social isolation, leading to worse long-term health trajectories.^8,9^ There are currently no FDA-approved therapeutics to address social deficits in ASD, largely due to a limited understanding of the neural mechanisms underlying these deficits. Thus, efforts to improve social behavior in ASD require a clearer understanding of the mechanisms driving social behavioral deficits.

The vast majority of ASD cases are attributable to genetic factors.^10^ The most prevalent genetic etiology is a complete deletion of the *SHANK3* gene, which encodes a postsynaptic scaffold protein essential for proper synaptic function.^11–15^ Our lab developed a *Shank3 exon 4-22* deletion (*Shank3*^Δ*e4–22*^) mutant mouse, which improves upon prior isoform-specific deletion models.^16^ *Shank3*^Δ*e4–22*^ mice show a social avoidance phenotype by spending less time in the social, mouse-paired chamber of a 3-chamber task, but the mechanisms underlying these deficits remain poorly understood.^16,17^ SHANK3 is highly expressed in the nucleus accumbens (NAc), a region known to regulate social behaviors.^18,19^ The NAc drives both approach and avoidance behaviors and integrates signals from many prosocial neural circuits, such as the amygdala and cortex, as well as receiving input of neural modulators such as serotonin, oxytocin, and dopamine, making it an attractive region for investigating the mechanisms underlying *Shank3*-mediated social behaviors.^20–30^ We and others have shown that SHANK3 protein expression in the NAc is necessary for social preference and social motivation, and that *Shank3* mutation causes atypical neuronal activity in the NAc and in NAc-associated circuits.^16,31–33^

The social motivation theory of autism posits that ASD-related mutations cause social deficits by impairing the neural processes that direct attention and motivation toward social cues ^34^, but this has not been well-investigated in *Shank3*^Δ*e4–22*^ mice. Therefore, we addressed this gap in knowledge by characterizing the function of the group of neurons that orchestrate social behavior, or social ensembles, in the NAc. Surprisingly, we demonstrate that while NAc social ensembles encode appetitive social cues in WTs, social ensembles encode avoidance in *Shank3*^Δ*e4–22*^ animals. We further demonstrate that NAc neurons in *Shank3*^Δ*e4–22*^ mice exhibit an enhanced modulatory response to social novelty, and suppressing social ensemble activity during social novelty prevents future social avoidance and restores social investigation. Taken together, our data show that *Shank3*^Δ*e4–22*^ mice have an enhanced NAc response to social novelty that drives social aversion, rather than an inability to direct attention or drive social motivation.

## Results

### *Shank3*^Δ*e4–22*^ NAc Neurons Are Hyperactive and Hypermodulated During Social Novelty in the 3-Chamber Task

To assess NAc neuronal recruitment by novel social exposure in *Shank3*^Δ*e4–22*^ and WT mice, we quantified c-Fos immunoreactivity after exploration of either a novel empty or social context (**Fig. S1A**). We identified a significant increase in c-Fos expression in the anterior NAc of WT mice paired with a social context, compared to WT mice paired with an empty context, while *Shank3*^Δ*e4–22*^ mice express similar levels of c-Fos in cells in the anterior and posterior NAc after exposure to both empty and social-paired contexts **(Fig. S1B, C)**. In the social context, we found that WT and *Shank3*^Δ*e4–22*^ mice exhibit similar social engagement with the novel target (**Fig. S1D, E)**. To ensure that the decreases in c-Fos expression were not due to anatomical discrepancies between genotypes, we also quantified neuronal nuclear protein (NeuN) expression **(Fig. S1F)** in the NAc Core **(Fig. S1G)** and NAc Shell **(Fig. S1H)** and found similar expression levels in WT and *Shank3*^Δ*e4–22*^ mice.

We next used one-photon miniature microscope calcium (miniscope) imaging during performance of the 3-chamber task in *Shank3*^Δ*e4–22*^ and WT mice to understand how single cells in the NAc respond to social investigation at time-scale relevant for behavior **(Fig. 1A)**. Importantly, neither miniscope recording **(Fig. S2 A-D)** nor single housing **(Fig. S2 E-H)** altered behavioral outcomes in the 3-chamber task. Implanted and singly housed WT mice, but not *Shank3*^Δ*e4–22*^ mice, show a significant preference for the social, mouse-paired chamber over the empty chamber **(Fig S2 A-H)**. We binned neuronal activity data into three epochs during the 3-chamber task: habituation, test, and the duration of the 1^st^ visit to the mouse-paired chamber (social novelty). NAc cells from both WT and *Shank3*^Δ*e4–22*^ mice show significant increases in event rate during habituation epochs, compared to test and social novelty epochs. **(Fig. 1B-D)**. Remarkably, NAc cells from *Shank3*^Δ*e4–22*^ mice show significantly higher event rates during the social novelty, test, and habituation epoch compared to WT mice **(Fig. 1D)**. *Shank3*^Δ*e4–22*^ mice showed a consistent increase in NAc activity during the social novelty epoch when we calculated the average event rate per mouse for each epoch **(Fig. 1E**), and when neuronal activity was measured by standardized trace activity (deltaf/ noise) per mouse **(Fig. S3 A, B)** and per cell **(Fig. S3 C**). Single cells in the NAc of WT and *Shank3*^Δ*e4–22*^ mice exhibit significant increases in positive correlation **(Fig. 1F)** and negative correlation **(Fig. 1G)** in the social novelty epoch compared to other epochs, indicating increased coordination of NAc cells during this epoch. *Shank3*^Δ*e4–22*^ mice show a significantly stronger negative correlation during the social novelty epoch than WTs **(Fig. 1G)**, suggesting more coordinated inhibitory regulation in this population.

**Figure 1:**
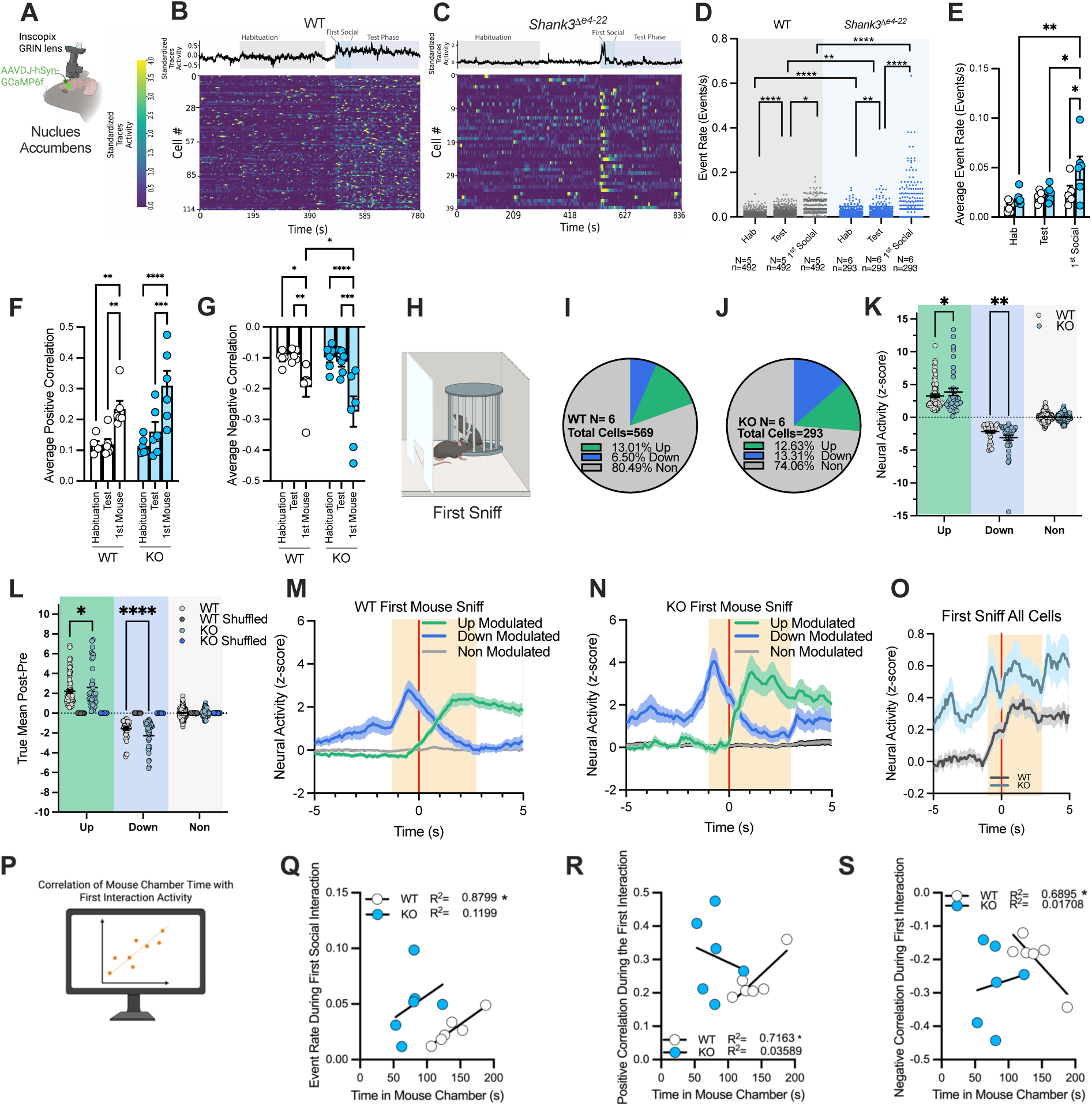
Single Cells in the NAc of Shank3^Δe4–22^ Mice Have Increased Activity and Modulation During the First Social Visit in the 3-Chamber Task. **A)** Single-cell imaging of NAc neurons was performed on WT and *Shank3*^Δ*e4–22*^ (KO) mice during the 3-chamber social interaction task. **B)** Representative heat map of neural activity from WT and **C)** KO mice during Habituation (grey, Hab), Test Phase (Purple), and First Social (Blue) phases of the task. **D)** Cells from KO mice have a significantly higher event rate than WTs in Habituation (****p<0.0001), Test (**p=0.0099), and First Social Epochs (****p<0.0001). WTs (****p<0.0001) and KOs (**p=0.0058) have higher event rates in test than habituation. WTs (*p=0.0367) and KOs (****p<0.0001) have higher event rates during 1^st^ social interaction. **E)** Event rate per cell was averaged per mouse. KO mice have significantly higher event rates during the 1st social epoch compared to WTs (*p=0.0120). Event rate was higher in the KO 1^st^ social epoch compared to habituation (*p=0.0160) and test epochs (**0.0039). **F)** Both WT and KO mice show significantly higher positive correlation of cells during the 1^st^ epoch compared to habituation (WT **p=0.0031, KO****p<0.0001) and test (WT **p=0.0028, KO ***p=0.0002) epochs. **G)** The NAc cells of KO mice show significantly more negative correlation in the 1^st^ social epoch compared to WTs (*p=0.04). KOs and WTs show the most negative correlation in the 1^st^ social epoch compared to the habitation (WT *p=0.0181, KO ****p<0.001) and test (WT **p=0.0092, KO ***p=0.0001) epochs (D-G: Mixed-effects analysis, Śidák’s multiple comparisons). **H-O)** Neuronal activity was analyzed between −1 and +3 seconds around the first time the mouse sniffs the mouse cup during the 3-chamber task to separate up- and down-modulated cells. **I)** WT mice have more cells captured than **J)** KOs during the 3-chamber social task, similar percentages of upmodulated cells between groups, but KO mice have double the percentage of downmodulated cells, compared to WTs. **K)** Both upmodulated (*p=0.0488) and downmodulated (**p=0.0019) NAc cells in KO mice are more responsive to the first sniff compared to WTs. **L)** The upmodulated (*p=0.0107) and downmodulated (p<****0.0001) cells in KO mice have significantly different means of activity (pre-post) of neuronal activity during the first sniff compared to WTs. (K,L 2-way ANOVA, Śidák’s multiple comparisons). **M)** Graphs showing neural activity by z-score of WT and **N)** cells separated by upmodulated, downmodulated, and non-modulated, or **O)** all cells grouped together. **P-S)** Total time spent in the mouse chamber was correlated with NAc activity metrics. WT, but not KO mice, showed a positive correlation with **Q)** event rate during 1^st^ social interaction (*p=0.0056), **R)** positive correlation of cell activity during 1^st^ social interaction (*p=0.03779), and **S)** negative correlation of cell activity during 1^st^ social interaction (*p=0.0407). (Q-S: Simple linear regression, R^2^ values in figure). KO= *Shank3*^Δ*e4–22*^.

To determine whether cellular hyperactivity was specific to the social novelty epoch, we also analyzed cellular activity during the 2^nd^ and the 1^st^ and 2^nd^ empty visits across genotypes **(Fig. S3D, E)**. NAc cells of *Shank3*^Δ*e4–22*^ mice were hyperactive in the social novelty epoch compared to the 2^nd^ social epoch and compared to the 1^st^ empty epoch **(Fig. S3E);** however, a caveat to this analysis is that the order of entries to the mouse and empty chambers is subject-specific, which may influence subsequent neural activity. To mitigate this, we also analyzed NAc cellular activity during the test epoch, excluding the social novelty epoch **(Fig. S3F-G)**. We found a significant decrease in neuronal activity in WT and *Shank3*^Δ*e4–22*^ mice following the first social visit (**Fig. S3F, G)**. Collectively, these data indicate that the social novelty epoch is a critical time point for NAc cellular response in both WTs and *Shank3*^Δ*e4–22*^ mice and that NAc cells are hyperactive in *Shank3*^Δ*e4–22*^ mice.

We next narrowed our analysis time window to immediately before (1s) and after (3s) the first sniff of the novel mouse cup in the 3-chamber task and clustered cells that were Up- and Down-modulated to this social novelty window **(Fig. 1H-N)**. Up- and Down-modulated NAc cells from *Shank3*^Δ*e4–22*^ mice exhibit significantly greater changes in neural activity in response to social novelty compared to WTs (**Fig. 1 K-N)**. We did not observe any changes in neural responses between genotypes when neural activity was analyzed immediately before and after the entry to the mouse chamber **(Fig. S4 A-H)**, indicating that the hyperactivity of *Shank3*^Δ*e4–22*^ NAc cells is specific to the investigation of a novel social target, and not the approach toward the novel social target.

To map neural activity back to behavior, we ran a regression analysis of NAc cellular activity during the social novelty epoch against total time spent in the mouse chamber across the entire 5-minute task **(Fig. 1P)**. In WT mice, there is a positive correlation between the event rate of NAc cells during the social novelty epoch and the amount of time mice spend in the mouse chamber, but this trend is absent in *Shank3*^Δ*e4–22*^ mice **(Fig. 1Q)**. We also ran a correlation analysis between average positive or negative cellular correlation during the social novelty epoch and time spent in the mouse chamber; again, we found a significant correlation in WT mice but not in *Shank3*^Δ*e4–22*^ mice **(Fig. 1R, S)**.

Of note, our imaging approach collected significantly more cells in the WT compared to *Shank3*^Δ*e4–22*^ mice **(Fig. 1I, J and S3 H)**; however, when we separated cells into Up-modulated, Down-modulated, and Non-responsive based on their activity during social novelty, we found similar cell counts of modulated neurons per mouse for each genotype **(Fig. S3 I)**. These data collectively provide critical clarifying temporal detail to our c-Fos analysis, which measured activity changes after the completion of the entire social dyadic task **(Fig. S1 B)**.

### NAc neurons of *Shank3*^Δ*e4–22*^ Mice Are Hypermodulated During Novel Nose-to-Nose Social Sniffing in the Social Dyadic Task

We next explored how NAc cells in *Shank3*^Δ*e4–22*^ and WT mice respond to a sex-matched juvenile mouse in a more naturalistic social dyadic task. Our social dyadic task was separated into 4 epochs: habituation, social habituation, social test, and social novelty. We hand-scored the behavioral data frame-by-frame during the social habituation and test phases using 12 behavioral markers that fell into naturalistic, social anxiety-like, social motivation, or pro-social behavioral categories **(Fig. S5 A-AA)**.^35^ *Shank3*^Δ*e4–22*^ and WT mice engaged in similar amounts of prosocial behavior during the test phase **(Fig. S5 Q-W)**, which we previously described.^17^ Here, we show that *Shank3*^Δ*e4–22*^ mice display significant increases in social anxiety-like behavior **(Fig S5 G-N)**, including tail rattling **(Fig S5 G, H)**, during the social habituation phase. Overall, our data reveal that *Shank3*^Δ*e4–22*^ mice shift away from neutral behaviors and toward social-anxiety-like behaviors predictive of social avoidance,^1^ while WT mice engage in greater exploration and social motivation during the social habituation phase **(Fig. S5 X-Y**). During the test phase, however, *Shank3*^Δ*e4–22*^ mice exhibit fewer behaviors across all domains **(Fig. S5 Z-AA)**.

We then quantified NAc neural event rates during the 4 epochs of the social dyadic task (**Fig**. **2A**). Single cells from the NAc of *Shank3*^Δ*e4–22*^ mice have increased event rates in the Habituation (Hab), Social Habituation (Social Hab), and Test (Free) Epochs compared to WT, but we found no differences in cellular activity during social novelty between genotypes (**Fig**. **2B**). These statistical differences were not found when we averaged event rate by mouse **(Fig. 2C)**. WT and *Shank3*^Δ*e4–22*^ mice both show stronger positive **(Fig. 2D)** and negative **(Fig. 2E)** correlation of neural activity during social novelty epoch; *Shank3*^Δ*e4–22*^ mice show a trend toward decreased positive correlation of cellular activity during a social novelty bout compared to WTs, but no differences were observed in negative correlation of activity.

**Figure 2:**
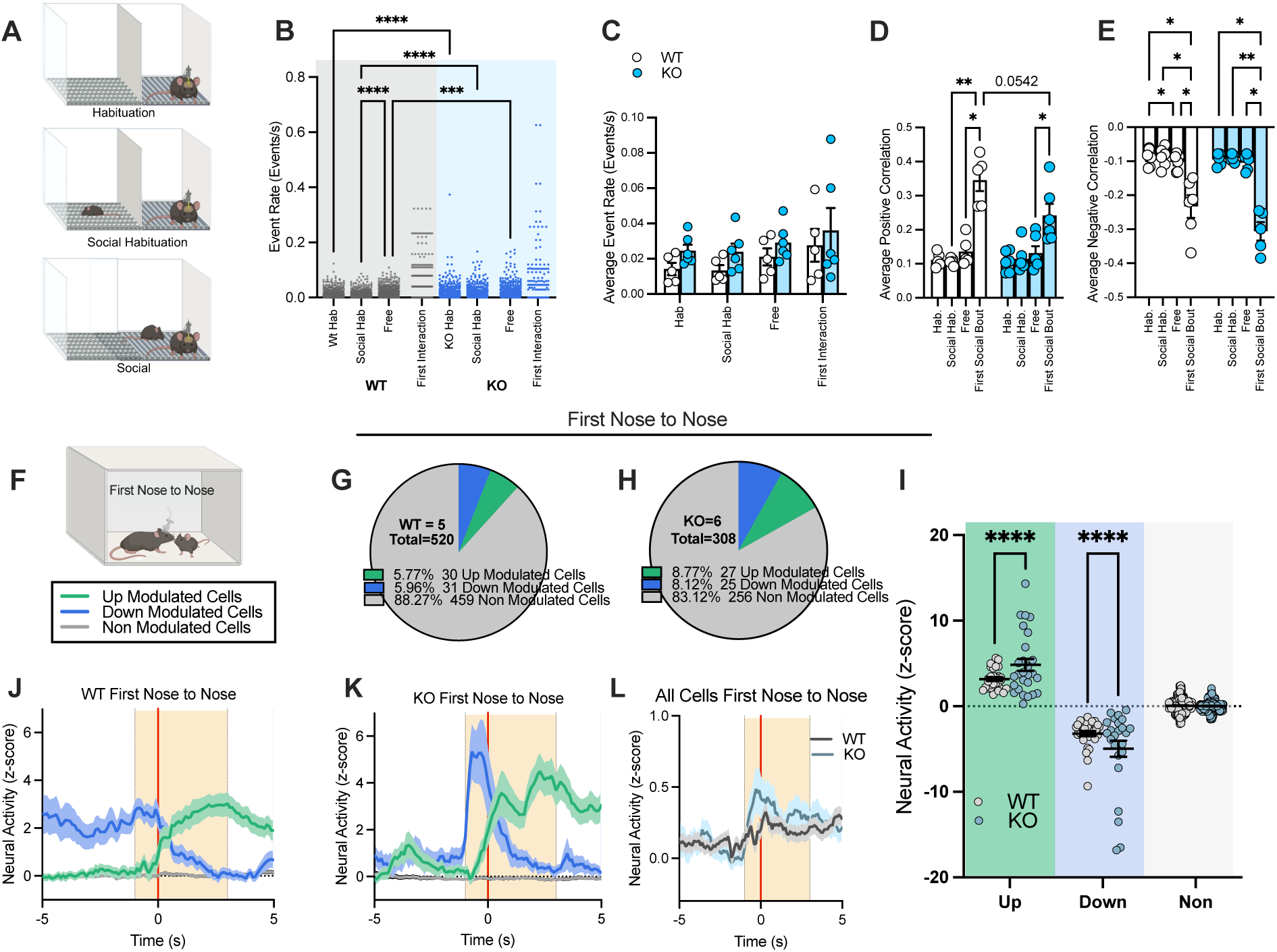
NAc Neurons of *Shank3*^Δ*e4–22*^ Mice Are Hypermodulated During First Nose-to-Nose Sniffing Bout. **A)** Single-cell NAc activity was recorded during a social dyadic task that was split into Habituation (Hab), Social Habituation (Social Hab), and Social Epochs (Free). **B)** Cells from *Shank3*^Δ*e4–22*^ (KO) mice have a higher event rate than WTs during the Social Habituation (****p<0.0001) and Test phase (***p=0.0001), but there are no differences during the first interaction epoch (Brown-Forsythe ANOVA with Dunnett’s T3 multiple comparisons test). **C)** There are no differences between epochs or genotypes when cells are averaged by mouse. **D)** Both WT (*p=0.0243) and KO (p=*0.0485) mice have an increase in positive correlation and **E)** negative correlation (WTs *p=0.0243, KOs p=*0.0485) of cell activity between the 1st social epochs and the test phase. **F)** Neuronal activity was analyzed between −1 and +3 seconds around the first nose-to-nose sniff of the free social dyadic task. **G)** WT mice have more cells captured than **H)** KOs during the social dyadic task. The percentage of upmodulated and downmodulated cells are similar between groups. **I)** The Upmodulated (Up) (****p<0.0001) and downmodulated (Down) (**p<0.0001) cells in KO mice have significantly different z-scores of neuronal activity during the first nose-to-nose event compared to WTs. **J)** Graphs showing neural activity by z-score of WT and **K)** cells separated by up (green), down (blue), and non-modulated cells (grey), or **L)** all cells grouped together. (B-E, I: Mixed-effects analysis, Śidák’s multiple comparisons).

While epoch analysis reveals critical information about how cells respond during each phase of this task, a caveat to this analysis is that the epoch for the first interaction is social behavior agnostic, as the social novelty epoch was defined as the first time the animals came in proximity to one another (∼ 5cm) until animals separated (∼ >5cm). Thus, we next directed our cellular analysis to examine the peri-events for prosocial behaviors: Nose-to-nose **(Fig. 2 F-I, Fig S6A-G**) and anogenital sniffing **(Fig S7A-N)**. During the first, novel nose-to-nose sniffing event, WT and *Shank3*^Δ*e4–22*^ mice recruit similar proportions of modulated neurons **(Fig. 2 G, H)**. *Shank3*^Δ*e4–22*^ mice have significantly increased neural responses in both Up-Modulated and Down-Modulated neurons **(Fig. 2 I-L**). When all nose-to-nose events were binned together, cells responded similarly between phenotypes **(Fig S6A-G),** further supporting the hypothesis that social novelty is a key time window for NAc social encoding of information in *Shank3*^Δ*e4–22*^ mice. NAc neural activity in *Shank3*^Δ*e4–22*^ and WT mice was similar during the first anogenital sniff (**Fig. S7 A-G**), suggesting nose-to-nose social interaction may be an important step in social novelty assessment and encoding of information.

### Chemogenetic Inhibition of the NAc Restores Social Preference in the 3-Chamber Social Task in *Shank3*^Δ*e4–22*^ Mice

Our *in vivo* imaging data revealed that although *Shank3*^Δ*e4–22*^ mice recruit fewer NAc cells during novel social investigation, the recruited cells are hyperresponsive to social stimuli. Therefore, we hypothesized that local inhibition of the NAc may restore NAc function and preference for social investigation in *Shank3*^Δ*e4–22*^ mice. To test this hypothesis, we injected *Shank3*^Δ*e4–22*^ mice with pan neuronal Gi-DREADD (AAV5-hSyn-hM4D(Gi)-mCherry) or AAV5-hSyn-mCherry control and injected 3mg/kg CNO 30 minutes before the 3-chamber social interaction task and free social task **(Fig. 3A-C)**. In the 3-chamber task **(Fig. 3D)**, *Shank3*^Δ*e4–22*^ mice injected with Gi-DREADDs showed a significant preference for the mouse chamber **(Fig. 3E)** and a preference to closely interact with the mouse **(Fig. 3F)**. However, the mCherry control group did not show a preference for either the mouse or the empty chamber. There were no differences between groups in distance traveled **(Fig. 3G)**. Furthermore, we found no differences between mCherry and Gi-DREADD-injected KO mice in either the free social dyadic task **(Fig. S8A-E)** or the open field task **(Fig. S8 F-H)**. Collectively, these data suggest that inhibition of NAc cells is sufficient to enhance social preference in *Shank3*^Δ*e4–22*^ mice.

**Figure 3:**
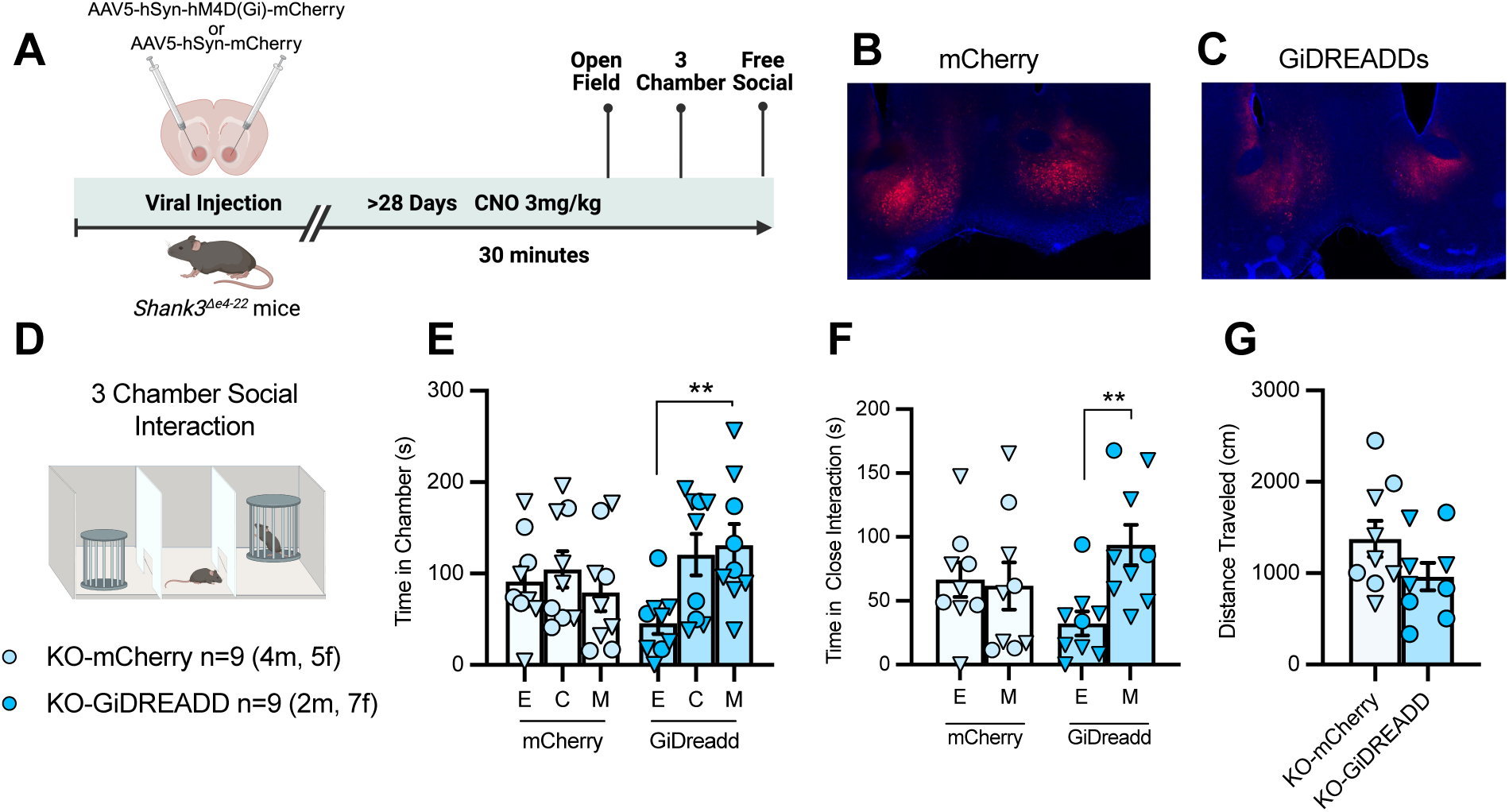
Chemogenetic Inhibition of the NAc of *Shank3*^Δ*e4–22*^ Mice Restores Social Preference in the 3 Chamber Social Task. **A)** Experimental Design. *Shank3*^Δ*e4–22*^ (KO) mice were bilaterally injected into the NAc with viruses encoding **B)** mCherry control or **C)** Gi-DREADD. CNO was injected 30 minutes before the **D)** 3-chamber social interaction task. **E)** Gi-DREADD injected mice have a significant preference for the mouse chamber (**p=0.0054), and **D)** close interaction (**p=0.0058) but not mCherry controls (Mixed-effects analysis, Śidák’s multiple comparisons). **E)** No differences were found in the distance traveled during the 3-chamber task. in the 3-chamber task (unpaired t-test).

### Inhibition of NAc Social Ensembles Restores Social Preference in *Shank3*^Δ*e4–22*^ Mice but Drives Social Preference Deficits in WTs

We next sought to identify and manipulate social ensembles, to determine their causal role in social behavior using <u>T</u>argeted <u>R</u>ecombination of <u>A</u>ctivated <u>P</u>opulations (TRAP) driven by the early-activation gene Arc (ArcTRAP).^36^ Cre-dependent halorhodopsin (DIO-NphR) and bilateral fiber optic implants were placed into the NAc of *Shank3*^Δ*e4–22*^ mice and WT mice that had been crossed with the *Arc^CreER^* and Ai14 mouse lines **(Fig. 4A)**. To capture social ensembles, mice were exposed to a novel conspecific in the social dyadic task and injected with the TRAP activator 4-hydroxytamoxifen to restrict TRAP expression to socially active cells **(Fig. S9A)**.^36,37^ We observed no differences in behavior during the social habituation phase **(Fig. S9 B-D)** or the social phase **(Fig. S9G, H)** during social ensemble capture between genotypes. Within the same cohorts, nonsocial ensembles were captured by placing mice into a new, empty arena without a social target.

**Figure 4:**
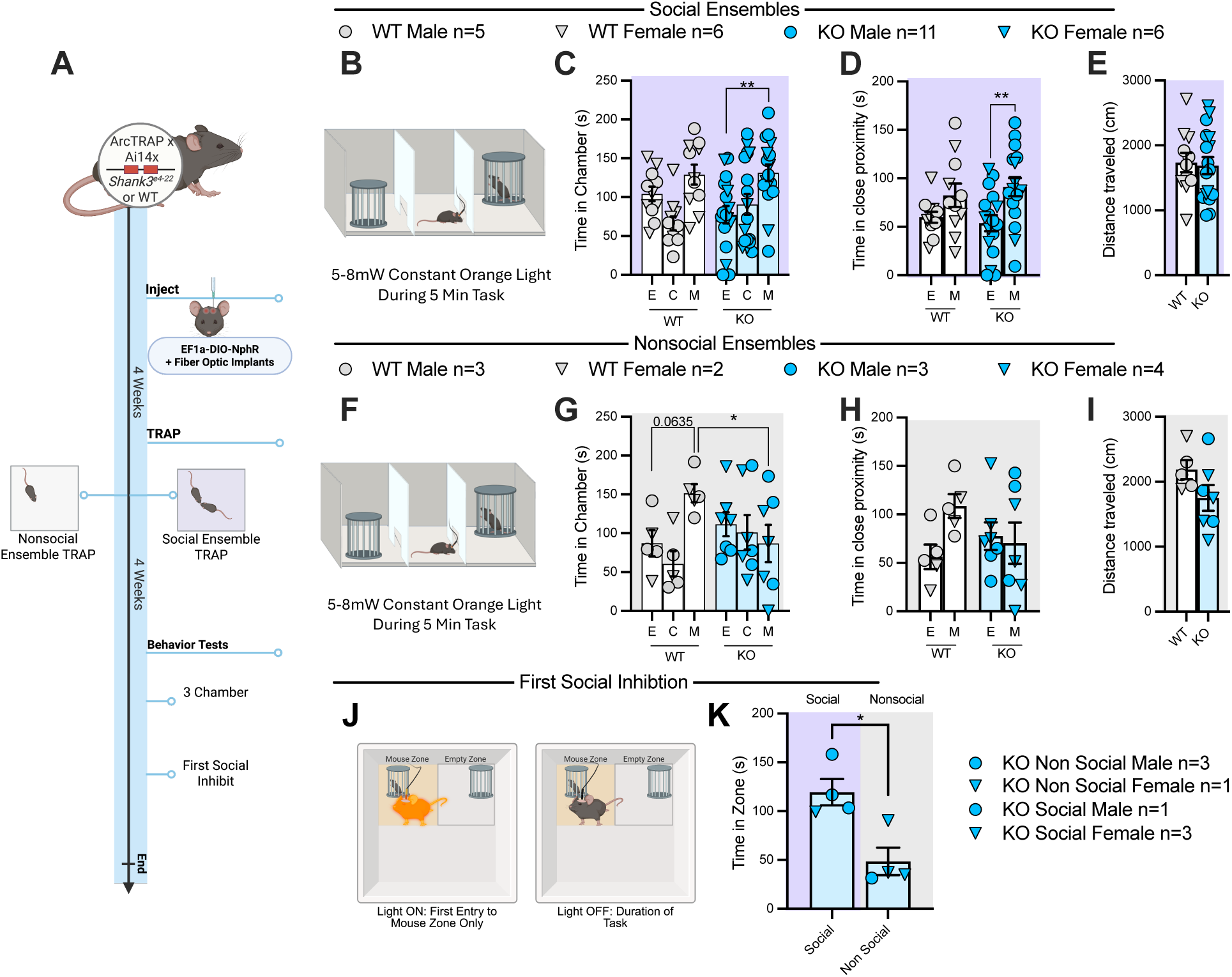
Inhibition of *Shank3*^Δ*e4–22*^ NAc social ensembles during the first social interaction is sufficient to rescue social deficits. **A)** Experimental Design. **B)** Social ensembles were inhibited during the entire duration of the 3-chamber task in WT and *Shank3*^Δ*e4–22*^ (KO) mice. **C)** While social ensembles are inhibited, KO mice show a preference for the mouse chamber (**p=0.0010) and **D)** Close interaction (within 5cm) zone (**p=0.0056) ((Mixed-effects analysis, Śidák’s multiple comparisons). **E)** No differences in distance traveled were observed. **G)** Nonsocial Ensembles were inhibited throughout the 3-chamber task. **G)** WTs have a trend toward preference of the mouse chamber and spent significantly more time in the mouse chamber compared to KO mice (*p=0.0331). **H)** No statistical differences were observed in close proximity time or **I)** distance traveled. **J)** Optogenetic stimulation was delivered only the first time that animals entered the social zone. **K)** Inhibiting social ensembles of KO mice during initial interaction significantly increases time spent in the social zone compared to inhibition of nonsocial ensembles (*p=0.0108). (C-D, G-H: Mixed-effects analysis, Śidák’s multiple comparisons, E, I, K: Unpaired t-test).

After 4 weeks, mice were placed in the 3-chamber social interaction task. During the test, orange light was delivered to the fiber optic implants to silence ensembles **(Fig. 4B)**. With social ensembles inhibited, we observed a significant preference for the mouse chamber **(Fig. 4C)** and close proximity to the social target **(Fig. 4D)** in *Shank3*^Δ*e4–22*^ mice, and a lack of chamber or close interaction preference in WTs. There were no differences in the distance traveled between genotypes **(Fig. 5I)**. On the other hand, inhibition of nonsocial ensembles in the 3-chamber task did not alter social investigation preference in WT or *Shank3*^Δ*e4–22*^ mice **(Fig. 4F)**. WT mice exhibited a trend toward preferring the mouse-paired chamber **(Fig. 4G)**, while *Shank3*^Δ*e4–22*^ mice spent significantly less time in the mouse chamber than WTs and showed no preference to either chamber **(Fig. 4G)**. Distance traveled did not differ between genotypes **(Fig. 4I)**.

Because our single-cell recordings showed that *Shank3*^Δ*e4–22*^ mice exhibited aberrant NAc activity predominantly during social novelty, we next tested the hypothesis that inhibiting social ensembles only during the first, novel social interaction would rescue social deficits in *Shank3*^Δ*e4–22*^ mice. Thus, *Shank3*^Δ*e4–22*^ mice were placed in an arena with a novel conspecific under a cup in one corner, and an empty cup in the other **(Fig. 4J)**. Orange optogenetic light was delivered only when animals first entered close proximity to the mouse cup and was turned off when they left the zone. This acute inhibition of social ensembles during social novelty was sufficient to significantly increase the time spent near the social conspecific throughout the duration of the task compared to acute inhibition of nonsocial ensembles **(Fig. 4K)**.

## Discussion

The present study shows that NAc social ensembles encode opposing social information in WT compared to ASD models. We demonstrate that NAc social ensembles are necessary for social preference in WTs but drive social avoidance in *Shank3*^Δ*e4–22*^ mice. Additionally, we show that *Shank3*^Δ*e4–22*^ mice exhibit stronger, more coordinated NAc cellular engagement during social novelty and suggest that *Shank3*^Δ*e4–22*^ social ensembles encode aversive social behavior. While social deficits in ASD have often been discussed in terms of reduced neural responsivity and diminished encoding of social motivation,^34^ our results offer an additional possibility that maladaptive hyperresponsivity drives social aversion in ASD models.

Our results show that NAc neuronal activity is highest during social novelty epochs in two well-established social tasks in WT and *Shank3*^Δ*e4–22*^ mice. These data are consistent with recent work showing that the NAc is a central hub for social novelty in WT mice.^38–43^ Our findings advance this framework by showing that aberrant hyperactivity of NAc cells in response to social novelty drives future social avoidance in *Shank3*^Δ*e4–22*^ mice. *Ex vivo* NAc hyperactivity, driven presynaptically, has previously been shown in *Shank3* mutant models, suggesting that the aberrant NAc activity reported here may be driven by maladaptive presynaptic inputs.^33^ Given that the NAc receives dopamine from the ventral tegmental area (VTA) during social novelty epochs,^39–41,44^ and removing *Shank3* from the VTA dampens VTA-NAc dopamine neuron firing,^45^ future studies should investigate whether alterations in VTA dopaminergic signaling contribute to the atypical social ensemble function and the hyperactive NAc activity during social novelty we identified in *Shank3*^Δ*e4–22*^ mice.

One possible explanation for the differences in the activity and function of social ensembles between *Shank3*^Δ*e4–22*^ and WT mice may be distinct variations in the cell types recruited. The majority (95%) of neurons in the NAc are GABAergic projection medium spiny neurons (MSNs), primarily expressing D1 or D2 dopamine receptors, with the remaining 5% being local interneurons.^22,46^ D1^+^/D2^+^ MSNs and interneurons regulate social play in rodents, and the activity of each of these cell types is dysregulated in *Shank3* mutants.^1,23,24,41,47–50^ It is therefore likely that the recruitment of D1^+^/D2^+^ MSNs and interneurons into social ensembles is different between *Shank3*^Δ*e4–22*^ and WT mice. In addition to better characterizing the ensemble, identifying cell type-specific functions within the NAc social ensembles will help address a major gap in our understanding of the mechanisms underlying social deficits in ASD. D2^+^ MSNs track prediction error, showing high activity when a stimulus is novel and decreasing as the stimulus becomes more predictable. Many studies suggest that activation of D2^+^ MSNs promotes social avoidance,^41,47,51^ but the role of NAc D2^+^ MSNs on social prediction error is not well understood. Future investigations are needed because disruptions in prediction errors are well-established in ASD and in many widely used ASD mouse models,^52–54^ and deficits in prediction errors may reveal a converging mechanism underlying social deficits in ASD. Our data support a model where *Shank3*^Δ*e4–22*^ mice overactivate and overrecruit D2^+^ MSNs in novel social environments, potentially driving social prediction errors and leading to social avoidance behaviors. Future research is necessary to verify this hypothesis.

The current work is not without limitations. First, ArcTRAP captures cells over a relatively long 12-hour window, which can lead to off-target cell capture. Although we single-housed mice at least a week before and after cell capture to restrict social activity, social isolation may also influence our results.^36^ New methods for neuronal capture with tighter time windows would enable reintegration into group housing, addressing this caveat.^55–58^ Second, further research is warranted to better understand how inhibiting social ensembles during social novelty improves social investigation in *Shank3*^Δ*e4–22*^ mice. Our study is limited to a 5-minute test with one novel conspecific. Using a longer test with additional novel and familiar conspecifics would clarify whether NAc social ensemble inhibition is needed for each novel conspecific and determine how long social ensemble inhibition improves social investigation in *Shank3*^Δ*e4–22*^ mice. Finally, although we use a highly valid and well-investigated *Shank3*^Δ*e4–22*^ model, characterizing NAc social ensembles in additional ASD mouse models is necessary to determine whether NAc social ensemble dysfunction is a converging mechanism underlying ASD social deficits.

Despite a 300% rise in autism diagnoses in the past two decades, treatment options remain limited due in part to a lack of understanding of the neural mechanisms of social deficits in ASD.^59^ In this study, we observed that silencing NAc social ensembles exclusively during social novelty increased social investigation in *Shank3*^Δ*e4–22*^ mice. These results mirror clinical findings showing that behavioral therapies for autism that provide support in novel social environments increase social salience, improve social competency, and reduce anxiety in future social engagements.^60–62^ Taken together with our work, we suggest that social novelty may be a critical temporal window for the incorrect encoding of social information in ASD, and research into pharmacological or circuit-based therapies targeted at social novelty is warranted.

## Methods

### Animals

*Shank3^De4–22^* heterozygous mice (Jax Stock #039524) were bred in the Jiang lab to generate *Shank3^De4–22−/−^* and WT littermate controls for all experiments. ArcTRAP (B6.129(Cg)-*Arc^tm1.1(cre/ERT2)Luo^*/J, Jax stock #021881)^36^ mice were crossed with Ai14 (B6.Cg-*Gt(ROSA)26Sor^tm14(CAG-tdTomato)Hze^*/J, Jax stock #007914)^63^ to visualize ensembles. This cross was ithen crossed with *Shank3^De4–22^* mice to generate male and female *Shank3^De4–22^* ^+/−^ x Ai14^+^ x ArcTRAP^+^ that were used as breeders to generate *Shank3^De4–22^*: ArcTRAP^+^:Ai14^+^ (*Shank3 ^De4–22^* TRAP) and WT:ArcTRAP^+^:Ai14^+^ (WT TRAP) used in experiments. WT targets for social behavior testing were C57BL/6J mice obtained directly from Jackson Laboratories, or the offspring of WT x WT crosses obtained from Jackson Laboratories.

Male and female mice were maintained on a 12-hour light/dark cycle, with lights on at 0700 hours, in a temperature- and humidity-controlled room. Mice were housed in Tecniplast GM500 cages with corncob bedding and one nestlet. An igloo was added in breeding cages. Food and water were available ad libitum. All cages were assigned to the same rack location in the animal facility, which remained consistent through all cohorts tested. All mice were on a C57/Bl/6J strain and ranged in age from 6 to 26 weeks. All experiments were conducted during the light phase.

Mice were kept group-housed by sex, unless where noted for specific experiments. Following stereotaxic surgery, miniscope-implanted animals and ArcTRAP mice were separated into single housing for the duration of the experiment. We also single-housed mice for at least 1 week prior to the 3-chamber social to ensure single housing did not change social investigation behaviors **(Fig S1)**.

All animals were monitored daily in their home cages by experimenters and Yale Animal Resources Center veterinary staff for activity, signs of stress and pain, weight, and postoperative recovery. Pertinent to these studies, animals were monitored during testing for aggression and skin lesions from over-grooming, which are common in *Shank3^De4–22^* mice.^16^ If any adverse health or activity-related events were observed, mice were treated by veterinary staff and removed from the study. We have complied with all relevant ethical regulations for animal use. Animal husbandry and all animal protocols received ethical approval by the Yale Animal Resources Center and the Institutional Animal Care and Use Committee at Yale.

### Viruses

*Shank3^De4–22^* and WT mice were injected with 300nL AAVDJ-hSyn-GCaMP6f (Diesseroth Vectors at University of North Carolina at Chapel Hill Vector Core) for imaging studies. 350nL of AAV5-Ef1a-DIO eNpHR 3.0-EYFP was bilaterally injected into *Shank3^De4–22^* and WT TRAP mice for ensemble-dependent silencing.^64^ pAAV-Ef1a-DIO eNpHR 3.0-EYFP was a gift from Karl Deisseroth (Addgene viral prep # 26966-AAV5; http://n2t.net/addgene:26966; RRID:Addgene_26966). 300-400nL of AAV5-hSyn-hM4D(Gi)-mCherry was used to silence activity or AAV5-hSyn-mCherry as a control. AAV5-hSyn-hM4D(Gi)-mCherry was a gift from Bryan Roth (Addgene viral prep # 50475-AAV5; http://n2t.net/addgene:50475; RRID:Addgene_50475). AAV-hSyn-mCherry was a gift from Karl Deisseroth (Addgene viral prep # 114472-AAV5); http://n2t.net/addgene:114472; RRID:Addgene 114472).

### Drugs

4-Hydroxytamoxifen (4-OHT) (50mg/kg; Sigma-Aldrich, H6278) was prepared in a 1:9 EtOH:peanut oil solution and injected I.P. at a volume of 10 µl/g bodyweight one hour after social interaction or novel chamber exposure.^37,65^ JZL-184 (8mg/kg; Cayman Chemical Company) was prepared in DMSO and injected at 1 µl/g body weight 2 hours before behavioral testing or 1 hour before miniscope recordings.^33,66,67^ Clozapine-N-Oxide (CNO) (3mg/kg, Hello Bio) was prepared in saline at a volume of 10 µl/g bodyweight and administered I.P. 30 minutes prior to experimental manipulation.

### Stereotaxic Surgery

Mice underwent stereotaxic surgery at 6-12 weeks of age, generally using a previously reported protocol.^17^ Animals were anaesthetized with 3-5% isoflurane and 1% oxygen, then administered Ethiqa XR 1.3mg/mL at 0.6mg/kg (S.Q., Covetrus) for preemptive analgesia. Mice were moved to a stereotaxic frame (RWD) and kept under anesthesia using 1% isoflurane, 1% oxygen delivered by nose cone. Before incision, mice were injected with bupivacaine (T.D., 1 mg/ml). The mouse’s fur was then shaved from the skull, and the skin was cleaned with 70% ethanol, iodine, and then fresh 70% ethanol. After incision, bregma and lambda were leveled within 0.1mm of each other. The skull was then scored with a scalpel blade and wiped clean with H2O2. A craniotomy was performed using a drill lowered into the skull until reaching the dura above the NAc. Dura was broken by hand with a syringe tip. Viruses were loaded into a NanoFil 10uL syringe with a 33-gauge blunt needle tip (World Precision Instruments). The needle was slowly moved into the NAc (anterior/posterior [AP]: −1.50mm; medial/lateral [ML]: +/− 0.75mm; dorsal/ventral [DV]: 4.50mm), paused for 1 minute, then moved to DV 4.45mm. The virus was automatically infused into the NAc at 100nL/ minute (Harvard Instruments). After infusion and a 10-minute wait, the needle was removed. For bilateral injection, the viral infusion protocol was repeated.

#### Optogenetic Implants

Following bilateral viral infusion, dual fiber-optic cannulas (Doric-Pitch 1.5mm, 200µM-NA0.37, 5mm length, flat tip, guiding socket top) were slowly lowered into the injection site using a 1.25mm stereotaxic cannula holder fitted with a guiding socket adapter (Doric). Pilot cohorts were implanted with two single fiber-optic cannulas. One was placed at a 0^°^ tilt (RWD, 5mm length, 200 µm core, 0.39 NA), and the other (7mm) was placed at a 10^°^ tilt to accommodate patch cable attachment to the headcap.

#### Miniscope Lens Implants

Following unilateral AAV-GcAMP injections, a guide needle (a 21.5g syringe needle that was clipped, filled, and sanded) was slowly lowered (approximately 600μm/min) to DV: 3.56mm (80% of final target). After a 5-minute wait, the guide needle was removed. Next a ProView™ Integrated Lens (0.6mm x 7.3mm, Inscopix) was inserted using an Inscopix Dummy Microscope to lower. The lens was lowered at approximately 400μm/min from a DV of 0 to 3.56mm, then at 100μm/min from 3.56 to 4.45mm.

#### Post Operative

All implants were secured to the skull using Metabond (Parkell). For viral and implanted animals, the incision site was sutured, and mice were given 1mL of saline S.Q. and 5mg/kg Carprofen I.P. Mice received 5mg/kg injections of carprofen every 24 hours for 72-hour recovery monitoring. Mice recovered for 3-4 weeks before any experiments.

#### Immunohistochemistry

##### Tissue

Tissue was collected using a previously reported protocol.^17^ In brief, mice were anesthetized using isoflurane in a bell jar and transcardially perfused with phosphate buffered saline (PBS, 10mL) followed by 4% paraformaldehyde in 0.1 phosphate buffer (PFA, 15-20mL). Brains were harvested and stored overnight in 4% PFA and transferred the following day to a 30% sucrose solution for at least 48 hours. Brains were cut at 80uM using a Leica CM3050 S cryostat or a Leica SM2000R microtome and slices were stored in an antifreeze, ethylene-glycol solution and stored at −20° until analysis. Tissue was then transferred to cell strainers and washed 4 times for 10 minutes each in fresh PBS. Next, slices were transferred to a PBS + 0.3% Triton-X 100 solution and washed for 60 minutes. Slices were then washed in a blocking buffer made of 3% normal donkey serum (Jackson immune research laboratories) in 0.3% Triton-X 100 PBS for 1 hour. Slices were then transferred to primary antibody solution overnight at room temperature. Anti c-Fos (Cell Signaling Technology #64555 1:1000), anti-GFP (abcam, ab13970, 1:1000), or anti NeuN (Sigma, MAB377 1:500), was diluted in blocking buffer. The next day, the slices were washed 4 times in fresh PBS for 10 minutes each, then incubated in Alexa 647 donkey anti-rabbit (Fisher Scientific A-21245) for c-Fos quantification, 1:1000 Alexa 594 Donkey Anti-Mouse (abcam, ab150108) for NeuN quantification, or 1:000 Alexa 647 donkey anti chicken at 1:1000 for 3 hours at room temperature. Slices were again washed 4 times for 10 minutes in fresh PBS and mounted onto slides using Fluoromount-G (EMS Acquisition) and cover slips were sealed with clear nail polish.

##### Viral and Implant Validation and Experiments

Tissue was transferred to cell strainers in a bath of fresh PBS for 10 minutes 4 times. Slices were then mounted onto charged slides and mounted usinf Dapi-Fluoromount-G and sealed with a coverslip and clear nail polish. Images were collected at 20x objective on a Zeiss 980 and manually reviewed for viral expression by a blind experimenter, which the lead author later unblinded.

##### Imaging

Images of slices were taken using a Zeiss 980 using 10x, 20x and 63x objective. Slices were compiled into a tiled image on one plane. For c-Fos and NeuN expression analysis, images were first deidentified so the experimenter could remain unbiased, and the NAc brain region was identified using a Mouse Brain Atlas (Paxinos and Franklin).

##### Analysis

Blinded experimenters then counted the number of c-Fos^+^, NeuN^+^, and DAPI^+^ cells by hand in Image J. We then calculated c-Fos per area using the image scale to determine the NAc area. c-Fos /mm^2^ was averaged across the left and right hemispheres. The data were later unblinded by the lead investigator for analysis.

#### TRAP

Test mice and social targets were handled for 3 days prior to TRAP. On the day before TRAP, mice received an I.P. injection of saline using a 23g needle in the room where behavior TRAP was to occur and were left to habituate to the room in their home cage for at least 1 hour. On the day of TRAP, all mice were first habituated to the room for at least an hour. Next, mice were put through the social dyadic (**see behavior methods**) task that included a 5 min habituation, 5 min social habituation, and test epochs. 1 hour after completion of the test phase, mice were injected with 50mg/kg 4-OHT I.P and retuned to home cage. If surgery was performed, TRAP occurred at least 4 weeks after.

#### Miniscope Recordings

Animals were connected to the Inscopix miniscope nVoke system. Recordings were collected in the inscopix data acquisition (DAQ) box. Recoding settings, including the focus, gain, and laser power, were established over 3 days prior to behavior testing. On test day, animal behavior was recorded with a camera connected to a USB-ethernet network adapter with a coaxial cable sync and trigger port (WhiteMatter), which was connected to the Inscopix DAQ box for direct TTL synchronization of behavior recordings with the miniscope recording data. The start of behavior recording was time-stamped in the calcium recording metadata to allow for frame-by-frame timing accuracy. Calcium and behavior videos were recorded at 30 frames per second.

##### Miniscope Preprocessing

Calcium and behavior videos were uploaded to the Inscopix Data Processing Software (IDPS). Videos were temporally cropped to remove frames prior to behavioral recording. Videos were spatially and temporally down-sampled by 2, then a spatial bandpass filter was applied from 0.005 to 0.5, and finally motion corrected using a region of interest around the lens. Then CNMFe was applied to detect cells using the following parameters in a parallel processing mode: cell diameter of 12, gaussian filter width of 1, minimum pixel correlation of 0.95, minimum peak to noise ratio of 12, a merging threshold of 0.5, patch overlap of 20, patch size 80, ring size factor of 1.4, background down-sampling of 2, and a closing kernel size of 1. Traces were then shown as deltaf / noise, and cells were manually accepted by analyzing the trace and comparing it with the processed calcium imaging video. Finally, event detection was performed in IDPS using the beginning of the trace as a reference, with an event threshold of 5.5 (in units of median absolute deviation (MAD) and defines a static threshold with respect to the input trace), and a decay constant of 0.2, while ignoring negative transients.

##### Miniscope Analysis

Processed calcium movies, event detection data, and cell sets were then uploaded to the Inscopix data exploration, analysis, and sharing (IDEAS) platform. Data was analyzed for epoch activity in the 3-chamber task using the compare neural activity across epochs function. Epochs were defined through manual review of the behavior video synchronized with the calcium movie in IDPS to determine the habituation, test, first social interaction, test after first social interaction, mouse visit 2, and empty visits 1 and 2 epoch times. Traces were rescaled using a standardization protocol in IDEAS ( in which calcium traces are standardized to have a mean of 0 and a standard deviation of 1. Events were not rescaled. These data were then combined by genotype using the “Combine and Compare Neural Activity Across Epochs” tool. Average trace activity and event rate for each cell during all epochs were plotted and analyzed both individually and averaged per mouse in PRISM 10. Positive and negative correlation data for each mouse across all epochs were also plotted and analyzed in Prism 10.

Data was analyzed from −1s before the event to 3s after the event and visualized from −5s before the event to 5s after. The following additional parameters were used: 1000 random shuffles, a significance threshold of 0.05, a seed of 0, and temporal down-sampling by a factor of 1. These data were then combined by genotype using the “Combine and Compare Peri-Event Analysis Data” tool with a significance threshold of 0.05, averaged by neurons, and with a tolerance of 0.5. Average total, population-level activity response to the event, average activity of upmodulated cells, average activity of downmodulated cells, and average activity of non-modulated cells were plotted from −5 to 5 around the event. Additionally, we plotted the z-scores for individual cells and grouped cells by modulation type. Finally, we plotted the cell counts for each modulation group by genotype. Data were visualized and compared across genotypes in PRISM10.

Next, neural activity was analyzed directly before and after specific events (e.g., first sniff, first social entrance, first empty sniff, first nose-to-nose, all nose-to-nose) using the peri-event analysis workflow in IDEAS. Data was analyzed from −1s before the event to 3s after the event and visualized from −5s before with event and 5s after. The following additional parameters were used: 1000 random shuffles, a significance threshold of 0.05, a seed of 0, and temporal downsampling by a factor of 1. These data were then combined by genotype using the “Combine and Compare Peri-Event Analysis Data” tool with a significance threshold of 0.05, averaged by neurons, and with a tolerance of 0.5. Average total, population-level activity response to the event, average activity of upmodulated cells, average activity of downmodulated cells, and average activity of non-modulated cells were plotted from −5 to 5 around the event. Additionally, we plotted the z-scored activity traces for individual cells and grouped cells by modulation type. Finally, we plotted the cell counts for each modulation group by genotype. Data were visualized and compared across genotypes in PRISM10.

#### Behavior Testing

Test and target mice were handled and tail-marked for 3 days before behavior testing and habituated to the testing room for at least 1 hour prior to the experiment. Before the 3-chamber test, target mice were acclimated to the pencil cups in their home cages for at least 3 days. All equipment was cleaned before, in between, and after each trial with 70% ethanol.

##### 3-Chamber Social Investigation Test

We followed previously reported metods.^17^ In brief, mice were placed in the center chamber of a 3 chamber apparatus (420mm x 565mm x 358mm box with 185mm x 420mm equal chambers made of 5mm thick, clear Plexiglas with two doors that were 120mm wide, 5mm thick, 358mm made by the Yale Machine Shop) that had two upside-down and empty wire pencil cups (Organize-it) placed in the outside chambers with full water bottles on top. After mice spent 5 minutes in the center chamber (habituation phase), an age and sex-matched target mouse was placed underneath the cup in one of the external chambers. The test mouse was then given 5 to 10 minutes to explore the entire arena. The target mouse location alternated across subsequent trials. Time spent in each chamber, time spent within 5cm of each cup (close interaction), and distance traveled were recorded. The experiment was conducted in 80-130 lux. The duration spent in the chambers was automatically scored using Noldus EthoVision XT (Leesburg, VA, USA). For miniscope recordings, the exact frames in which mice engaged in behaviors were hand-scored in the Inscopix Data Processing Software (IDPS, Bruker) to ensure that the behavioral and calcium videos were synchronized at the frame level.

##### Juvenile Social Dyadic Task

Test mice were placed in one side of a white, opaque plexiglass box (30cm × 30cm) fitted with clear inserts to create two equal chambers joined by a door (Yale Machine Shop). After 5 minutes in one chamber of the apparatus (Habituation), a sex-matched juvenile (< or =5 week-old mice) was placed in the opposite chamber for 5 minutes (Social Habituation). After another 5 minutes, the door was opened, and mice were permitted to freely interact for 10 minutes. Automated analysis of total Body Contact time was performed using Noldus EthoVision XT. Videos were also analyzed post-hoc by an experimenter by hand to capture the exact video frames at which behaviors were initiated, including: grooming, open exploration, freezing, tail rattling, social vigilance, flee/withdrawal, social reaction, approach, nose-to-nose sniffing, anogenital sniffing, side sniffing, and time spent in the same chamber.

#### Optogenetic Experiments

Animals were habituated to cables by being connected to a split fiber-optic patch cord for 10 minutes per day for 3 days prior to the experiments. (200µm, 0.37nA, 2m length, with a GS adapter at 1.5 separation. Pilot studies used a 200µm, 0.37nA, 1m length patch cord that was connected to implants via a 1.25mm zirconia sleeve; Doric). On test day, the optogenetic light source (LISER LD450, Doric) was connected to a fiber optic patch cord (480µm, 0.63nA, 1m length mono patch cord, Doric) that was connected to a 1×1 fiber optic rotary joint (doric). This rotary joint was then connected to the fiber-optic patch cables, which in turn connected to the animals’ implants. Orange, 593nm light was delivered using a 593 nm LISER Bandpass filter (Doric). Light was delivered at 5-8mW for all experiments.

### Exclusion criteria

Mice were excluded according to *a priori* criteria. Mice were removed if the implant and viral targeting or viral expression did not meet standard criteria. The targeted site was identified using microscopy to visualize the implant track and the presence of the viral-encoded fluorescent marker and by anti-GFP immunohistochemistry. Animals were excluded from all data sets if the viral expression was not expressed in all targeted regions. Genotypes were also confirmed post hoc, and mice were removed if genotype results were not confirmed. Finally, mice were excluded if they showed adverse health events after surgery, and 5 mice were excluded after an adverse reaction to 4-OHT.

### Statistics And Reproducibility

Statistical analysis was conducted using Prism 10 (GraphPad). Statistical significance was set at α = 0.05, and a *P* value less than 0.05 was considered significant. Generally, unpaired t-tests were used to compare 1 variable between two groups, One-way ANOVA was used to compare 1 variable with 3 groups with Tukey’s multiple comparisons test to correct for multiple comparisons. Mixed-Effects Analysis was used to analyze multiple variables between multiple groups with Šídák’s multiple comparisons test to correct for multiple comparisons. We used a Brown-Forsythe test to compare the standard deviations between groups; when the standard deviations were significantly different, a Welch’s correction or Welch’s ANOVA was used, with a Dunnet’s multiple comparisons test. Mann-Whitney nonparametric test was used to compare discrete data (i.e.: number of entries). Full statistical details and raw data values are in Supplementary Data 1 and sample size, statistical tests, and parameters are indicated in the figure legends. Data was not analyzed by sex, as we previously reported no sex differences in the behavioral phenotypes of *Shank3^De4–22^* mice^16^. Sample size was determined by referencing previous published literature from our group and others that use the same methods.^16,17,33,68^ Error bars represent mean ± standard error of the mean (SEM), with individual plot points overlaid. An annotation of * p<0.05, **p<0.01, *** p<0.001, and ****p<0.0001 is used in the figures. Detailed statistical results, including the tests and multiple comparisons used for all results, can be found in Supplementary Data 1.

## Competing interests

Y-H.J. is a scientific co-founder of Couragene Inc. and medical director of CourageAS Therapeutics, but this study is unrelated to his role. All other authors declare no competing interests.

## Acknowledgments

This research was supported by the NIH grants of HD088007 MH104316, MH098114, MH117289, HD087795 to YHJ. OMF was supported by the Eunice Kennedy Shriver National Institute of Child Health and Human Development F32HD106666. This work was also supported by the Kavli Institute for Neuroscience at Yale University, Kavli Postdoctoral Award for Collaborative Excellence to OMF. Some figures were created in BioRender (Folkes, O. (2026). The authors would like to sincerely thank Dr. Jing Liang-Guallapa, Dr. Sinem Erisken, and Dr. Qing Liu for their consultation on the miniscope experimental design, technical support, and data preprocessing.

