## Supplemental Data for "Social Novelty Recruits a Dysfunctional Nucleus Accumbens Ensemble That Drives Social Avoidance in a *Shank3^−/−^* Autism Model"

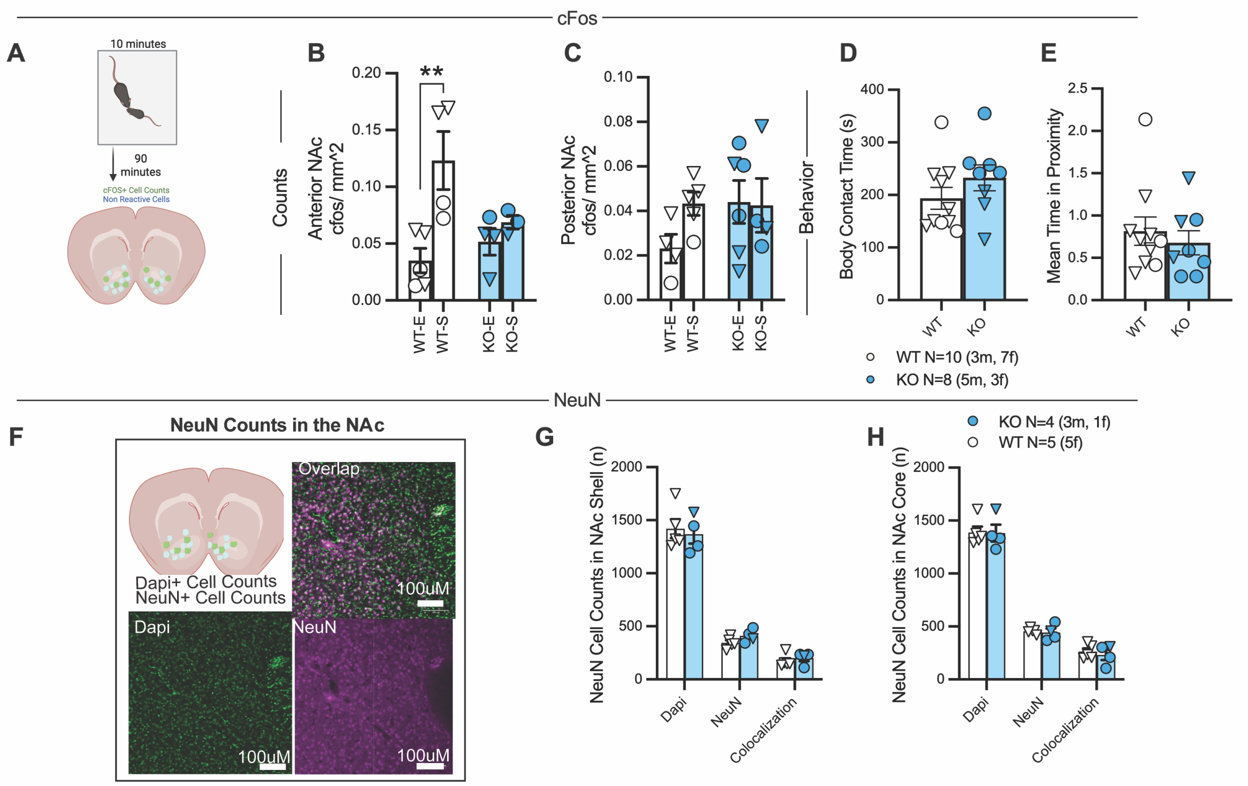


**Supplement 1: *Shank3*$^{\Delta e4\text{-}22}$ mice have fewer socially active NAc cells, but no change in neuronal density. A)** Experimental Schema. WT and *Shank3*$^{\Delta e4\text{-}22}$ mice were briefly exposed to a novel sex-matched mouse or an empty chamber. **B)** In the anterior NAc (1.7mm to 1.22mm from bregma) there are significantly more c-Fos$^+$ cells per mm$^2$ in WT mice after social exposure than WT after empty (**p=0.0027), but not *Shank3*$^{\Delta e4\text{-}22}$ mice. (Two-way ANOVA with Śidák's multiple comparisons). **C)** In the posterior NAc (1.20mm to 0.7mm from bregma) there are no significant differences between groups. **D-E)** During free social interaction, WT and *Shank3*$^{\Delta e4\text{-}22}$ mice used for c-Fos$^+$ analysis show similar time spent in **D)** body contact and **E)** time in proximity to the social target (Unpaired t-tests). **F)** Representative histology images showing Dapi (Green) and NeuN (Purple) staining. **G)** There were no differences in DAPI or NeuN$^+$ cell counts between WT and *Shank3*$^{\Delta e4\text{-}22}$ mice groups in the NAc Core or **J)** NAc Shell (Two-way ANOVA with Śidák's multiple comparisons). KO= *Shank3*$^{\Delta e4\text{-}22}$

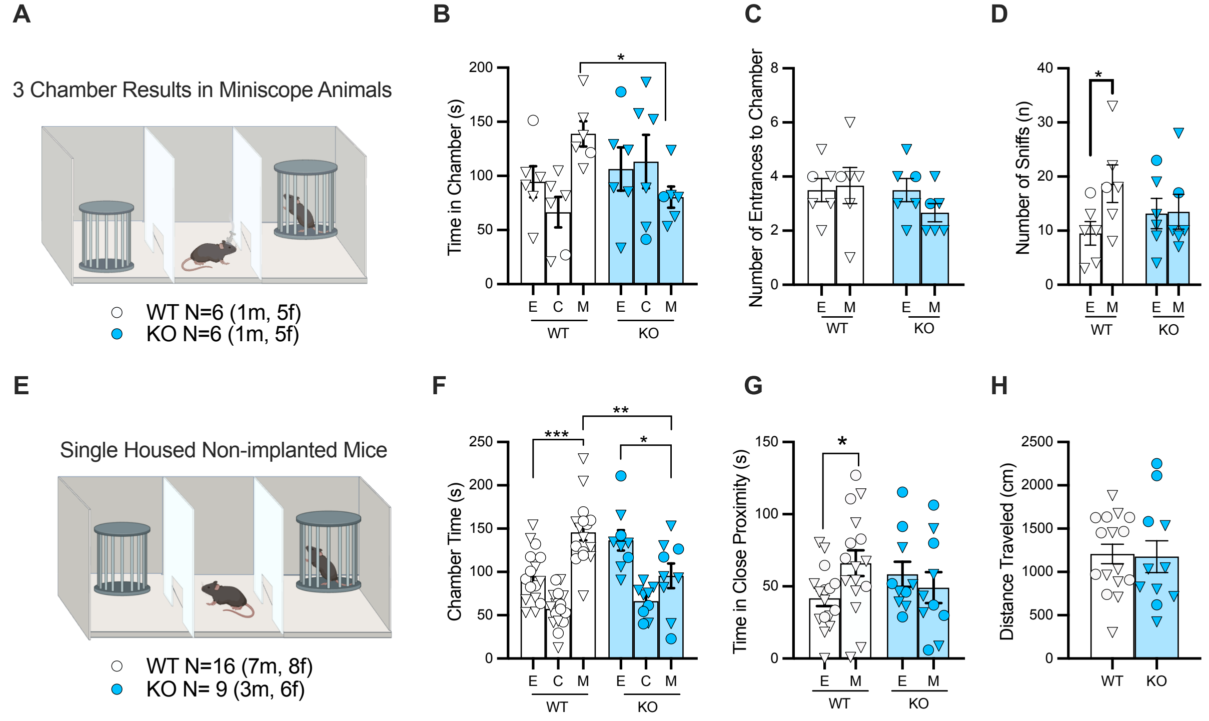


**Supplement 2: *Shank3*$^{\Delta e4\text{-}22}$ Mice Have Pervasive Deficits in Social Investigation. A-D)** Animals implanted with miniscope in 3 chamber task. **A)** Experimental Schema. **B)** *Shank3*$^{\Delta e4\text{-}22}$ mice spend significantly less time in the mouse-paired chamber compared to WTs (*p=0.0197). **C)** WT and *Shank3*$^{\Delta e4\text{-}22}$ mice enter the mouse chamber at similar rates, but **D)** WT mice sniff the mouse-paired cup significantly more than the empty cup (*p=0.0493) (Mixed-effects analysis Śidák's multiple comparisons). **E-H)** Social behavior in the 3-chamber task is unchanged after single housing. **E)** Experimental Schema. **F)** WT mice prefer the mouse chamber (***p=0.0005), while *Shank3*$^{\Delta e4\text{-}22}$ mice prefer the empty chamber (*p=0.0382). *Shank3*$^{\Delta e4\text{-}22}$ mice spend less time in the mouse chamber compared to WTs (**p=0.0032). **G)** WT mice prefer close interaction (within 5cm) (*p=0.0294) to the mouse cup, but not *Shank3*$^{\Delta e4\text{-}22}$ mice (Mixed-effects analysis Śidák's multiple comparisons). **H)** There were no changes in distance traveled during the 3-chamber task.

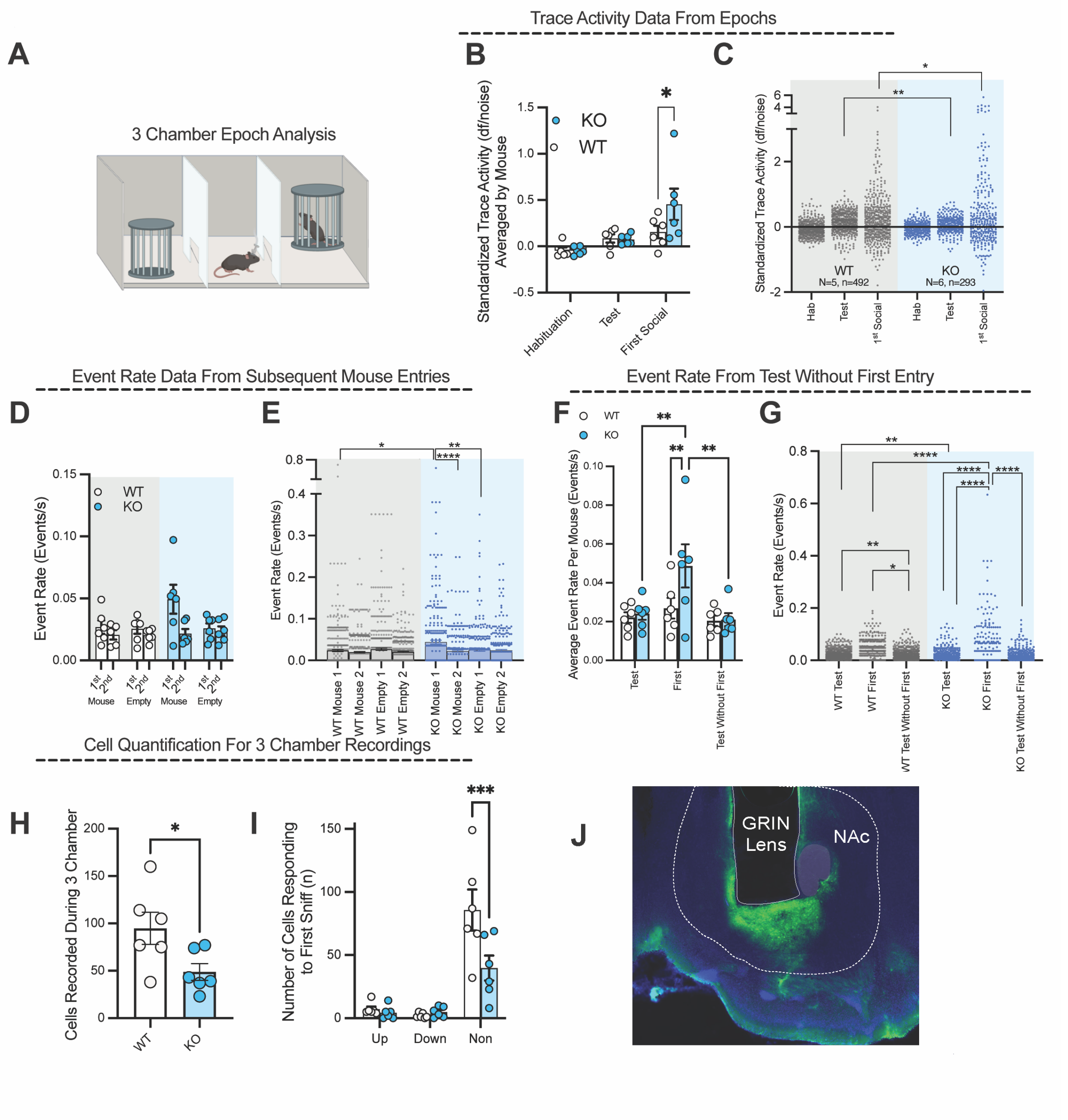


**Supplement 3: Extended Analysis of NAc cells during 3-chamber social interaction task. A)** Experimental Schema. **B)** Standardized trace activity averaged by mouse (*p=0.0387) and **C)** individually show that KO mice have higher NAc cellular activity during the 1$^{st}$ social epoch (*p=0.0381), but lower activity during the test epoch (**p=0.0032) compared to WTs. **D-E)** Event rate was similar between genotypes for 2$^{nd}$ mouse and 1$^{st}$ or 2$^{nd}$ empty epochs across **D)** mice and **E)** single cells. The event rate of KO NAc cells was significantly higher in the 1$^{st}$ mouse epoch compared to the 2$^{nd}$ mouse visit (****p<0.001), empty chamber (**p=0.0035), and WT 1$^{st}$ mouse epoch (*p=0.0417) as previously described. **F-G)** The test epoch was analyzed after the 1$^{st}$ entry to remove the initial activity spike. **F)** The event rate in KO mice during the 1$^{st}$ social epoch (first) is significantly higher than WTs (**p=0.0098), the test phase (**p=0.0055), and the test phase with the 1$^{st}$ social epoch removed (**p=0.0019). **G)** When the 1$^{st}$ social epoch was removed from the test epoch, there were no differences between genotypes. Event rate of NAc cells from KO mice is significantly higher in the 1$^{st}$ social epoch compared to KO test epoch (****p<0.0001), test without first (****p<0.001), and WT first social epoch (****p<0.001). **H)** There were significantly fewer cells captured in KO mice compared to WTs during recording (*p=0.0455, Mann-

Whitney test), but no differences in the number of **I)** Up or Down-Modulated cells between genotypes during the first social sniff. There were significantly fewer non-modulated (non) cells captured in KO recordings compared to WTs (***p=0.0008). (B-G, H: Mixed-effects analysis Śidák's multiple comparisons)

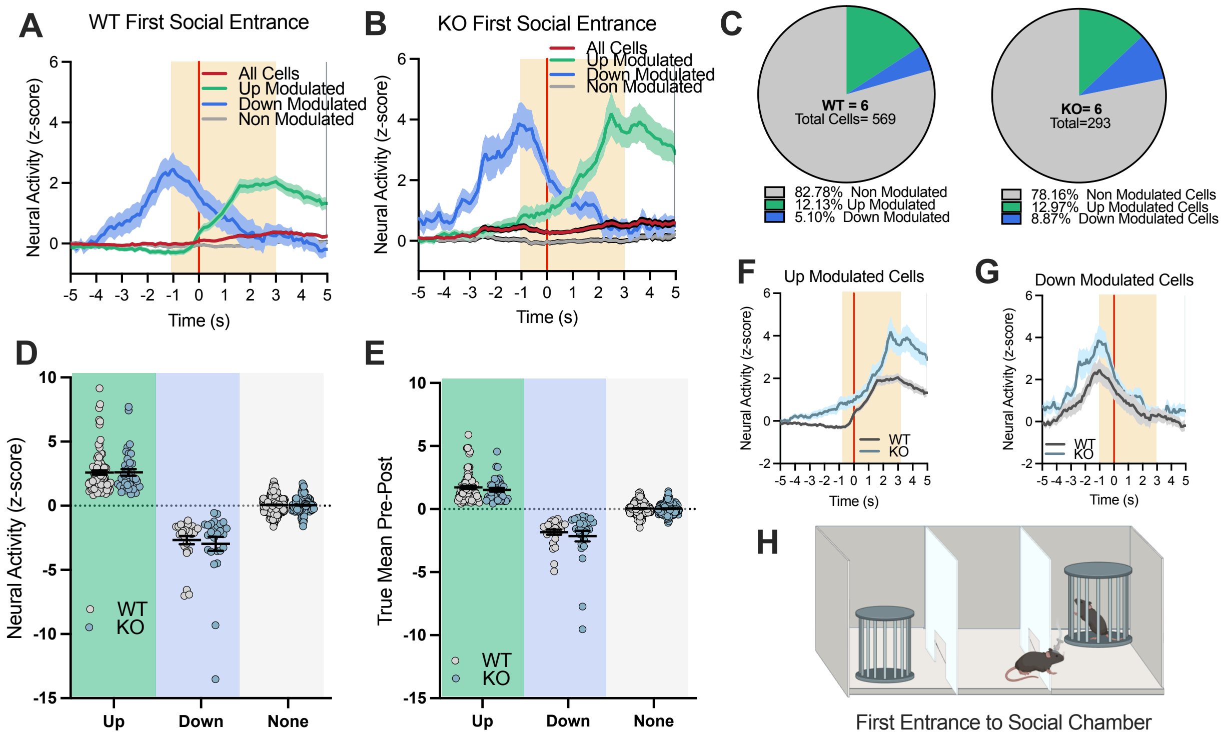


**Supplement 4: There are no differences in NAc cellular activity between genotypes when animals first enter the social chamber A)** WT and **B)** *Shank3*$^{\Delta e4\text{-}22}$ (KO) neural activity traces were analyzed -1 to +3 seconds when animals first crossed into the mouse chamber. There were no differences in the **C)** percentages of modulated cells, **D)** neural activity or **E)** true mean of **F)** Up or **G)** Down modulated cells during the **H)** First entry to the social chamber. D, E: 2way ANOVA, Śidák's multiple comparisons.

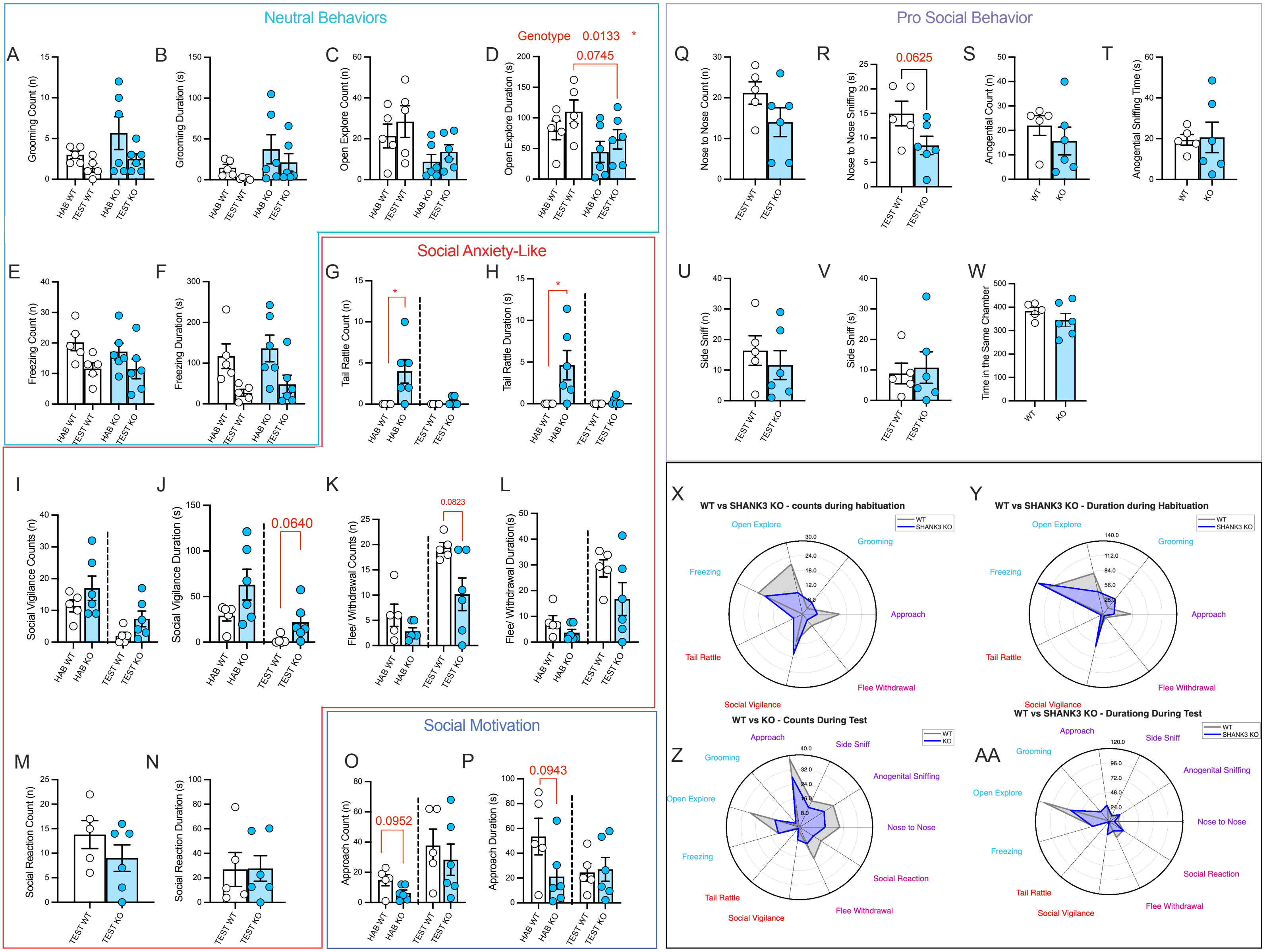


**Supplement 5: Frame-by-frame behavioral analysis of social habitation and test phases of the free social task in miniscope-implanted mice. A-F)** WT and *Shank3*$^{\Delta e4\text{-}22}$ (KO) mice exhibit similar neutral behaviors during both phases of the free social task. **G-N)** Social-anxiety like behaviors were modestly increased in KO mice including increased **G)** tail rattle counts (*p=0.0152, Mann-Whitney test), **H)** and tail rattle duration (*p=0.0466, Welch's t-test) during the habituation phase. **O-N)** There were no significant changes in social motivation, or **Q-W)** Pro-social behaviors between genotypes. **X-AA)** Behavior data summarized into spider plots. **X)** Counts and **Y)** Duration of behaviors during habituation show over-representation of social-anxiety-like behaviors in KO mice and increased neutral behavior in WTs. **Z)** KO mice engage in less behavior counts and **AA)** duration overall, but follow similar distribution patterns during the test phase .

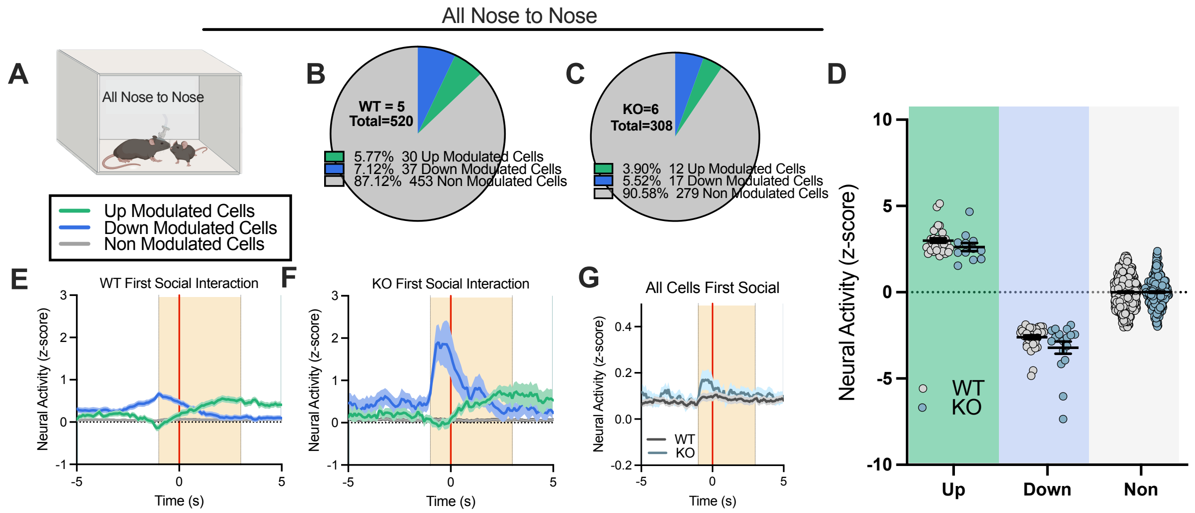


**Supplement 6: There are no differences between genotypes when all nose-to-nose events were analyzed. A)** All nose-to-nose events during the test phase of the social dyadic task were aligned. In both **B)** WT and **C)** *Shank3*$^{\Delta e4-22}$ (KO) mice, few cells responded to all nose-to-nose events, and **D-G)** there were no differences between genotypes across the activity of cell-responder types. (D: Mixed-effects analysis, Śidák's multiple comparisons)

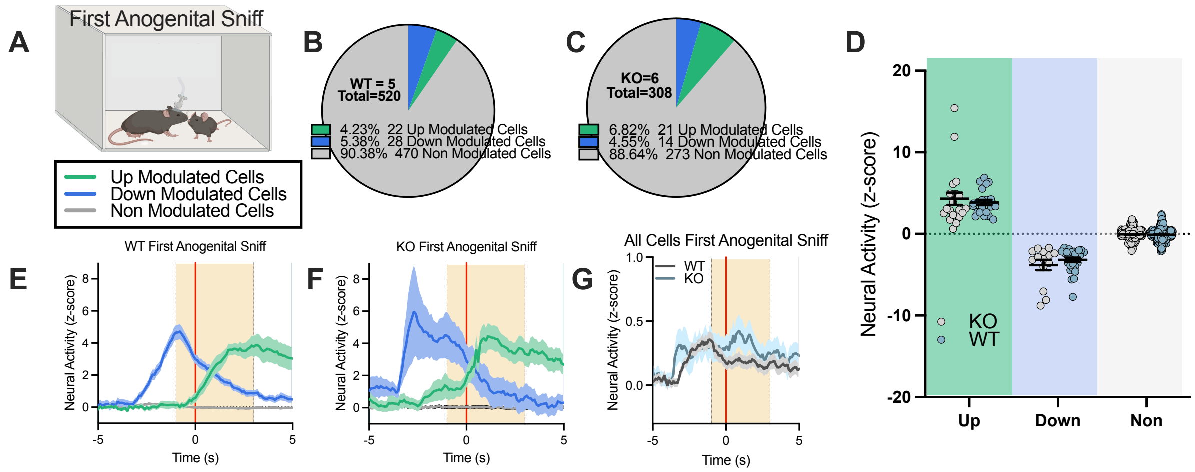


**Supplement 7: There are no differences in cell activity between genotypes for first or all anogenital sniffing events. A-G)** Cellular activity was analyzed -1 to 3+ seconds around the first anogenital sniffing event to reveal NAc cells from **B)** WT and **C)** WT mice have similar recruitment and **D-G)** activity in Up, Down, and Non-modulated cells. **A-G)** Cellular activity was analyzed -1 to 3+ seconds around the first anogenital sniffing event to reveal NAc cells from **B)** WT and **C)** WT mice have similar recruitment and **D-G)** activity in Up, Down, and Non-modulated cells. D: Mixed-effects analysis, Śidák's multiple comparisons)

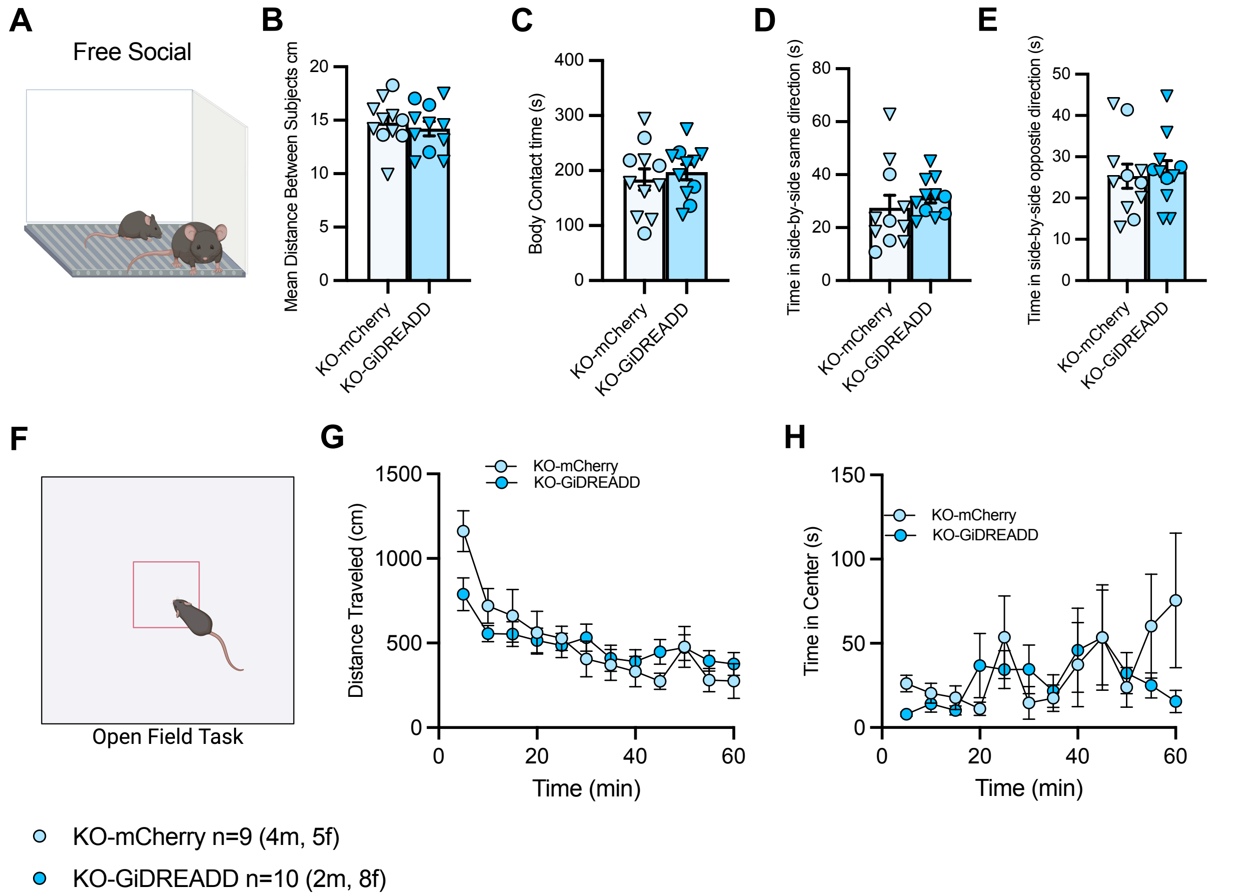


**Supplement 8: Chemogenetic Inhibition of the NAc of *Shank3*$^{\Delta e4\text{-}22}$ Does not alter Behavior During Free Social Dyadic or Open Field Task A-E)** There were no differences between mCherry and Gi-DREADDs-injected *Shank3*$^{\Delta e4\text{-}22}$ (KO) mice in **B)** distance between subjects, **C)** body contact time, or **D, E)** Time in side-by-side contact in the free social task (Unpaired t-test). **F-H** There were no differences between mCherry and Gi-DREADDs-injected KO mice in **G)** distance traveled or **H)** time spent in the center during the open field task.

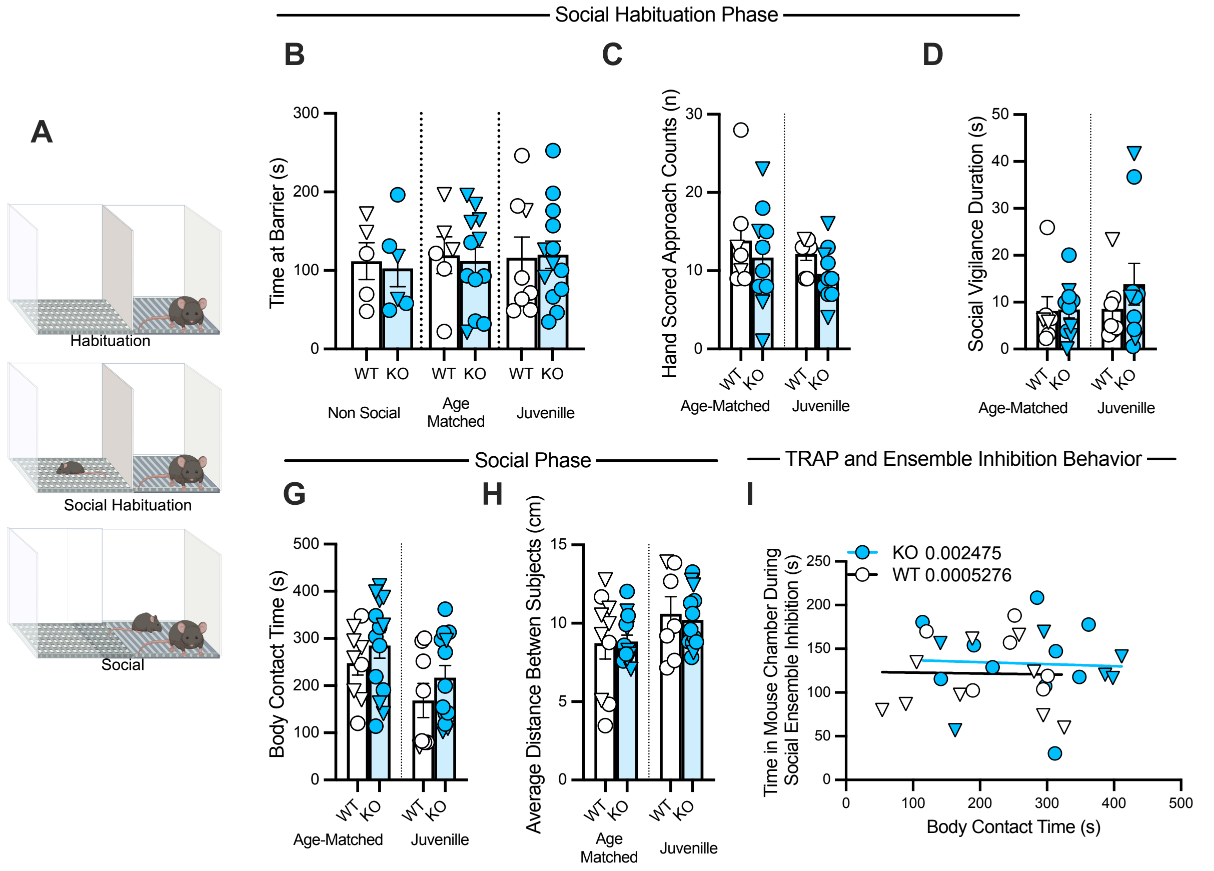


**Supplement 9: WT and *Shank3*$^{\Delta e4\text{-}22}$ mice exhibit similar social behaviors during the social dyadic task used to capture social ensembles. A)** WT and *Shank3*$^{\Delta e4\text{-}22}$ (KO) mice went through a 3-phase social dyadic task to capture social ensembles. There were no differences in **B)** time spent at the barrier, **C)** number of approaches to the barrier, or **D)** social vigilance behavior between genotypes or age of target during the social habituation phase. **G)** During the social test phase, there was an effect of target age on body contact time (*p=0.0159) but no differences in the **H)** average distance between subjects. (B-H: Two-way ANOVA, Śidák's multiple comparisons test) **I)** The time spent in body contact with the target during capture of social ensembles was not correlated with the effect of inhibition of social ensembles in either genotype (Simple linear regression).
